# Single-cell Mayo Map (*scMayoMap*): an easy-to-use tool for cell type annotation in single-cell RNA-sequencing data analysis

**DOI:** 10.1101/2023.05.03.538463

**Authors:** Lu Yang, Yan Er Ng, Haipeng Sun, Ying Li, Lucas C.S. Chini, Nathan K. LeBrasseur, Jun Chen, Xu Zhang

## Abstract

Single-cell RNA-sequencing (scRNA-seq) has become a widely used tool for both basic and translational biomedical research. In scRNA-seq data analysis, cell type annotation is an essential but challenging step. In the past few years, several annotation tools have been developed. These methods require either labeled training/reference datasets, which are not always available, or a list of predefined cell subset markers, which are subject to biases. Thus, a user-friendly and precise annotation tool is still critically needed. We curated a comprehensive cell marker database named *scMayoMapDatabase* and developed a companion R package *scMayoMap*, an easy-to-use single cell annotation tool, to provide fast and accurate cell type annotation. The effectiveness of *scMayoMap* was demonstrated in 48 independent scRNA-seq datasets across different platforms and tissues. *scMayoMap* performs better than the currently available annotation tools on all the datasets tested. Additionally, the *scMayoMapDatabase* can be integrated with other tools and further improve their performance. *scMayoMap* and *scMayoMapDatabase* will help investigators to define the cell types in their scRNA-seq data in a streamlined and user-friendly way.

## 1 Introduction

Tissues are constructed of diverse cell types that support highly specific functions in multicellular species. The development of single-cell sequencing (scRNA-seq) technologies has enhanced our capability to understand the molecular profiles of these individual cells and enabled us to study the heterogeneous cellular composition of complex tissues in the context of development, aging, health, and disease [1, 2]. In scRNA-seq data analysis, cell type annotation is a critical step and can be done manually with sufficient knowledge but is labor intensive and time-consuming. To achieve automatic cell annotation, computational tools have been developed to annotate either cells or cell clusters. Cell-annotation methods such as *SingleCellNet [3], SingleR, scmap [4]*, and *Azimuth [5]* assign cell identity to individual cells based on a pre-annotated scRNA-seq dataset as a reference or training dataset. However, accurate annotated reference data is not always available. Other tools, such as *SCINA* [6] and *CellAssign* [7], assign cell types based on known marker genes but they could be prone to the biases of the markers used.

In some cases, an annotation tool is accompanied by a cell marker database. For example, *scCATCH [8]* built a reference database “*CellMatch”* combining databases *CellMarker [9], Mouse Cell Atlas (MCA)* [10], *CancerSEA [11]*, and *the CD Marker Handbook [12]. SCSA* [13] integrated databases from *CellMarker* and *CancerSEA*. Some of these databases (e.g., *MCA*) were derived from differential expression analysis of scRNA-seq data. Other expert-curated databases (e.g., PanglaoDB) were manually curated from thousands of published studies. The annotation accuracy strongly depends on the informativeness and comprehensiveness of the marker gene database. To date, existing databases do not have extensive coverage in tissue types and cell types with good specificity. These obstacles can be challenging for investigators who are new to the scRNA-seq field or have limited background knowledge on the tissue and cell types involved.

To achieve a better cell type annotation outcome, we integrated and refined the available scRNA-seq annotation databases, originating a new database named *scMayoMapDatabase*. In parallel, we developed a new annotation tool, *scMayoMap*. Together with its own database *scMayoMap* provides easy, fast, and accurate cell type annotation without the need to provide cell type markers or pre-annotated single cell reference datasets. We envision *scMayoMap* will be of considerable utility for the scientific community.

## 2 Methods

### 2.1 Construction of *scMayoMapDatabase*

The data used in *scMayoMapDatabase* (including tissue, cell type, and marker genes, Table S1) was retrieved from literature and public databases including *PanglaoDB* [14], *Azimuth* [5], *CellMarker [9], CellMatch* [8], and *MCA [10]*. Marker genes from undefined tissues and cancer cells were not included. Individual tissues and cell types of each organ system were manually reviewed by experts in each organ system. The name of each tissue and cell type from different sources were standardized, and the corresponding cell markers were integrated. Both mouse and human genes were included in the database. Unofficial gene names were revised based on *Mouse Genome Informatics* [15] and *GeneCards* [16].

### 2.2 Cluster annotation process

The central algorithm for cell type annotation is based on the hypothesis that a good marker gene will be highly expressed in the cell type but not in unrelated cells. Based on this hypothesis, we designed the following steps to annotate cell clusters (Fig. 1).

**Figure 1.**
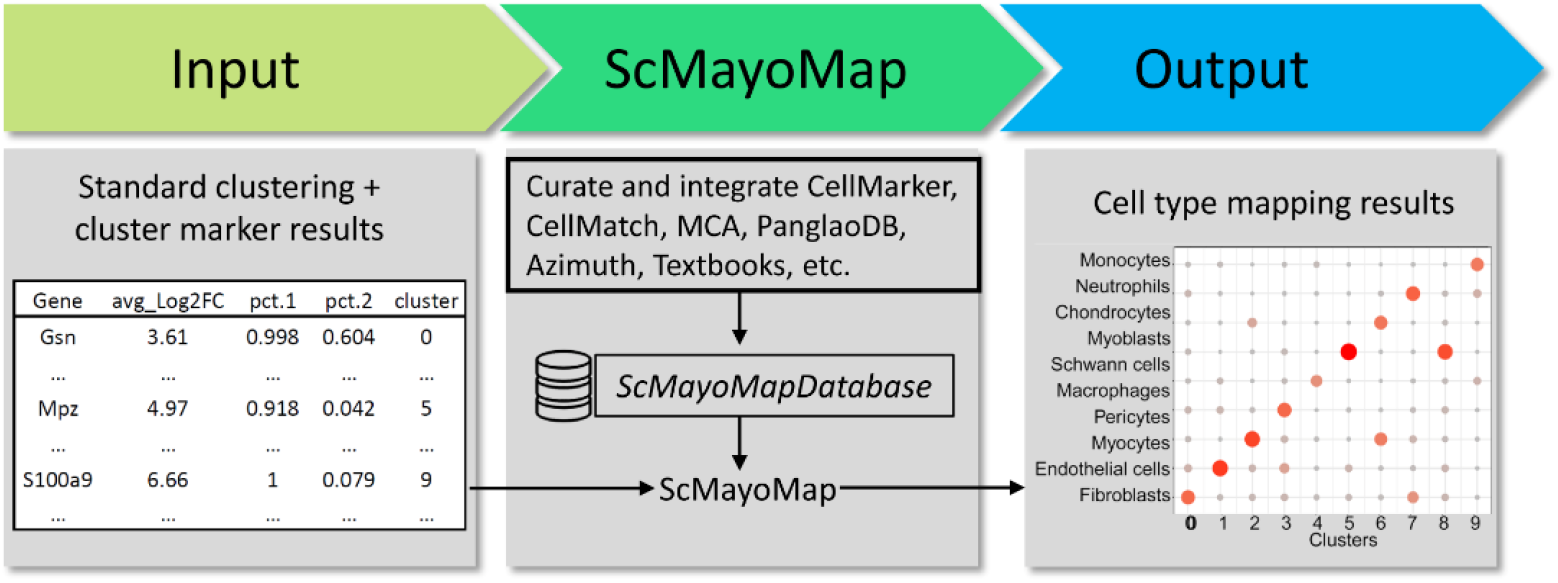
scMayoMap workflow. scMayoMap takes the standard cluster marker gene list as input and returns the cell type prediction results in a plot and the mapped gene list.

#### (1) Identification of potential marker genes for each cluster

Differential expression analysis is performed to find the marker genes for all *K* clusters. Gene expression in each cluster *k* (1 ≤ *k* ≤ *K*) is compared to the combination of all other clusters. This process can be easily done by applying the *FindAllMarkers* function in the *Seurat* package [5].

#### (2) Cluster annotation with potential marker genes

*scMayoMap* uses marker genes with percentage of cells expressing the marker gene greater than 0.25 and adjusted p-value less than 0.05. For cluster *k*, marker gene *i*, we produce a composite expression score 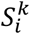 synthesizing both the fold change and the prevalence change: 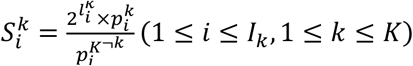, where 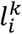denotes the average log_2_fold change of marker gene *i* in cluster *k* compared to other clusters. 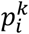 denotes the percentage of cells in cluster *k* where the gene *i* is detected, and 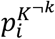 denotes the percentage of cells in all other clusters where the gene *i* is detected. The logarithm of 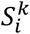 is essentially the sum of the log fold change and log prevalence change. After that, *scMayoMap* matches the marker genes in cluster *kk* to tissue-specific cell markers from *scMayoMapDatabase* and produces a score for each potential cell type *c* (1 ≤ *c* ≤ *C*) and cluster *k* (1 ≤*k* ≤ *K*), which is determined as follows:

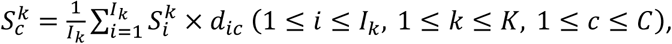

where *d*_*ic*_ is a binary indicator indicating the absence (*d*_*ic*_ = 0) and presence (*d*_*ic*_ =1) of the marker gene*i* in the cell type *c*.

To be more robust, *scMayoMap* allows assignment of multiple cell types to the same cluster if their evidence is similar. Specifically, the predicted cell types for cluster *k* are determined by the top *n* (1 ≤ *n* ≤ *C*) of 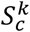, where *nn* is determined by finding the jump point in the cumulative variance of 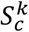 (see details at Supplementary Materials).

#### (3) Visualization of the prediction results

*scMayoMap*.*plot* function takes the cell-type prediction scores for each cluster as the input and produces a dot plot showing the predicted cell types for each cluster.

### 2.3 Experimental datasets

We retrieved 48 scRNA-seq datasets across different platforms and tissues from public resources (see details in Table S2). These datasets were generated by six different platforms, including Smart-seq2, CEL-Seq2, 10x Chromium, Drop-Seq, Seq-Well, and inDrops protocols [17]. Eighteen tissue types were included for the analysis, including blood, bladder, brain, adipose tissue, heart, kidney, gastrointestinal tract, muscle, liver, lung, mammary gland, bone marrow, pancreas, skin, spleen, thymus, trachea, and tongue [18, 19]. The cell types annotated in the source datasets were used as the benchmarks.

### 2.3 Comparison between different annotation tools

Four scRNA-seq-specific annotation tools, including two cell annotation methods (*scmap [4]* and *singleR* [20]) and two cluster annotation tools (*scCATCH* [8] and *SCSA* [13]) were compared. All methods were run with their default settings or based on the provided vignette describing the procedures. Briefly, for *scmap*, we used the scmap-cluster projection strategy that map the experimental dataset to “pbmcsca” reference dataset from SeuratData package. For *SingleR*, the *HumanPrimaryCellAtlasData* was used as the reference and “label.main” was specified as the prediction level. For *scCATCH*, tissue and species were specified for the analysis. For *SCSA*, the default database, whole.db, was used with input file generated by FindMarkers function of *Seurat*. To keep method evaluation on the same level, we used the proportion of cells that were correctly annotated as a metric. For cluster annotation methods, the predicted cell type with the maximum prediction score was chosen as the final predicted cell type. The annotation accuracy is the percentage of cells with correct cell type labels based on the source datasets. Cell types in source datasets with ambiguous cell type names, such as pp, MHC class II, PSC, co-expression, immune other and unclassified, were excluded from evaluation.

## 3. Results

### 3.1 Construction of *scMayoMapDatabase*

*scMayoMapDatabase* covers 340 cell types from 28 tissues and contains a total of 26,487 cell markers for both human and mouse (Fig. 2A and Table S1). It contains data for 12 organ systems including the integumentary system (skin), musculoskeletal system (bone, muscle), nervous system (brain), cardiovascular system (heart), circulatory system (blood), respiratory system (lung), digestive system (tooth, esophagus, liver, pancreas, stomach, small intestine, large intestine), urinary system (bladder, kidney), reproductive system (breast, ovary, placenta, uterus, mammary gland, embryo, testis, prostate), lymphatic system (bone marrow, thymus, spleen), endocrine system (adipose tissue), and visual system (eyes). The annotation level of the collected cell types for most tissues remains at the first level of the hierarchy. Cell markers for each cell type range from a minimum of 4 up to 300, with a mean of 43.8. Quality of the existing databases are partly compromised by tissues with very few cell types and cell types with very few marker genes. These limitations increase the noise for cell type annotation. Most collections in *scMayoMapDatabse* passed the quality control with more than 6 cell types per tissue and more than 4 marker genes per cell type. The Azimuth database has a good data quality, but the coverage of tissue types is relatively low (10 tissue types) (Fig. 2B).

**Figure 2.**
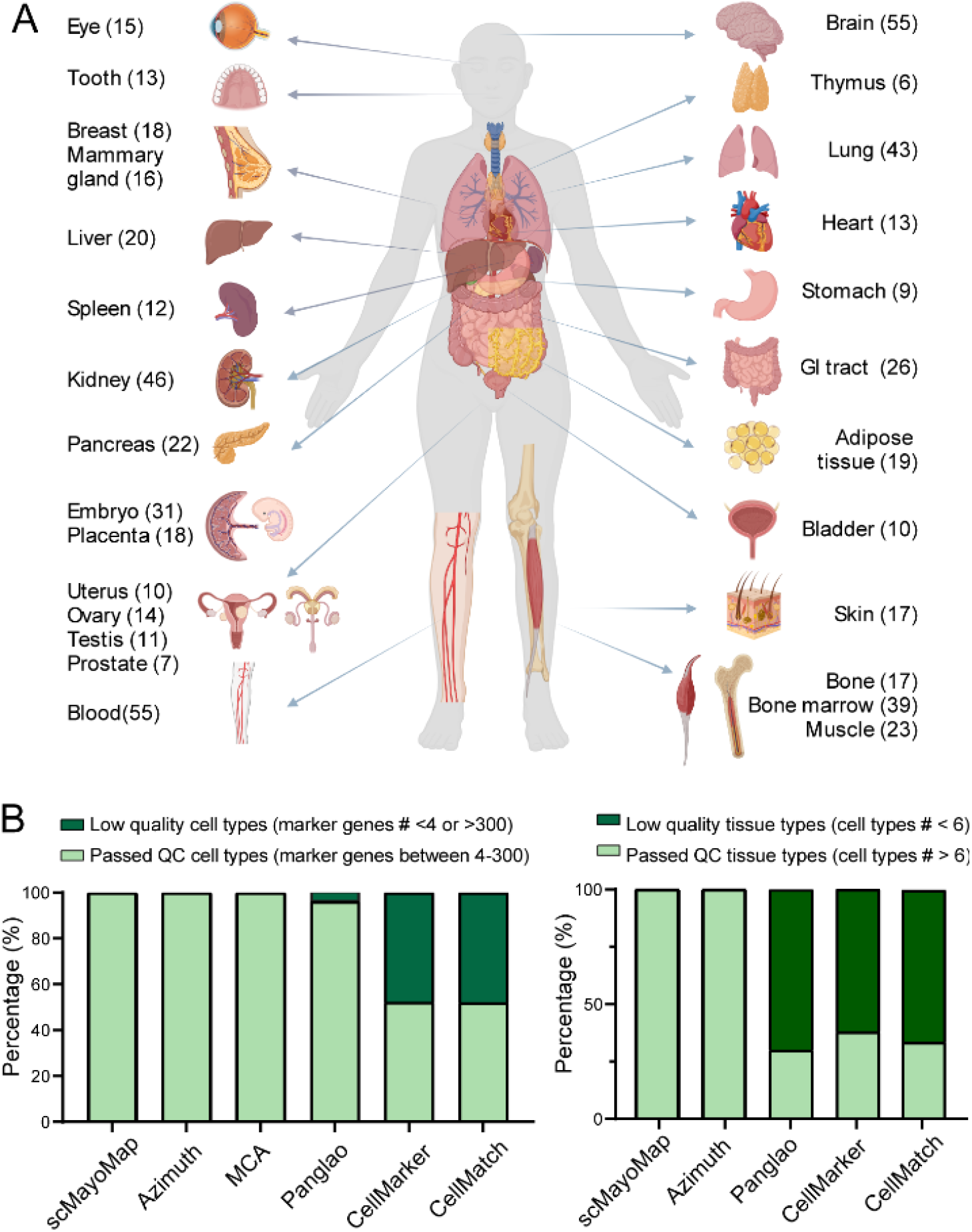
Summary of the scMayoMapDatabase. (A) The scMayoMapDatabase contains 28 tissues and 340 cell types for human and mouse. The number of cell types within each tissue was labeled in the parentheses. (B) Comparison of scMayoMapDatabase to other public databases. Left panel shows the cell type information and the right panel shows the tissue information in each database.

### 3.2. scMayoMap excelled in evaluation across multiple tissues and platforms

To evaluate the annotation accuracy of *scMayoMap* on a broad range of tissues, we retrieved data from the *Tabula Muris* [21] and literature [18, 22-26], covering 48 different scRNA-seq datasets, 18 different tissues, and 6 different scRNA-seq platforms (Table S2). *scMayoMap* annotated cell types from different tissues with high accuracy (Fig. 3 and Table S3). Specifically, it correctly annotated all clusters in 33 of 48 scRNA-seq datasets. *Tabula Muris* contains two different datasets that either uses droplet method or Smartseq2 method. For droplet-based datasets, *scMayoMap* successfully identified all cell types within tissues including bladder, heart, muscle, mammary gland, thymus, and spleen (Figure 3A). For Smartseq2-based datasets, more tissues were annotated accurately including bladder, brain, fat, limb, liver, pancreas, spleen, thymus, and trachea (Figure 3B). When we compared the different pancreas datasets, *scMayoMap* successfully annotated all datasets with 100% accuracy. *scMayoMap* correctly identified α, β, δ, ε, and γ cells of the islet, as well as acinar cells, pancreatic progenitor cells, ductal cells, and endothelial cells, macrophages, mast cells, stellate cells, Schwann cells, B cells, and T cells (Fig. 3C).

**Figure 3.**
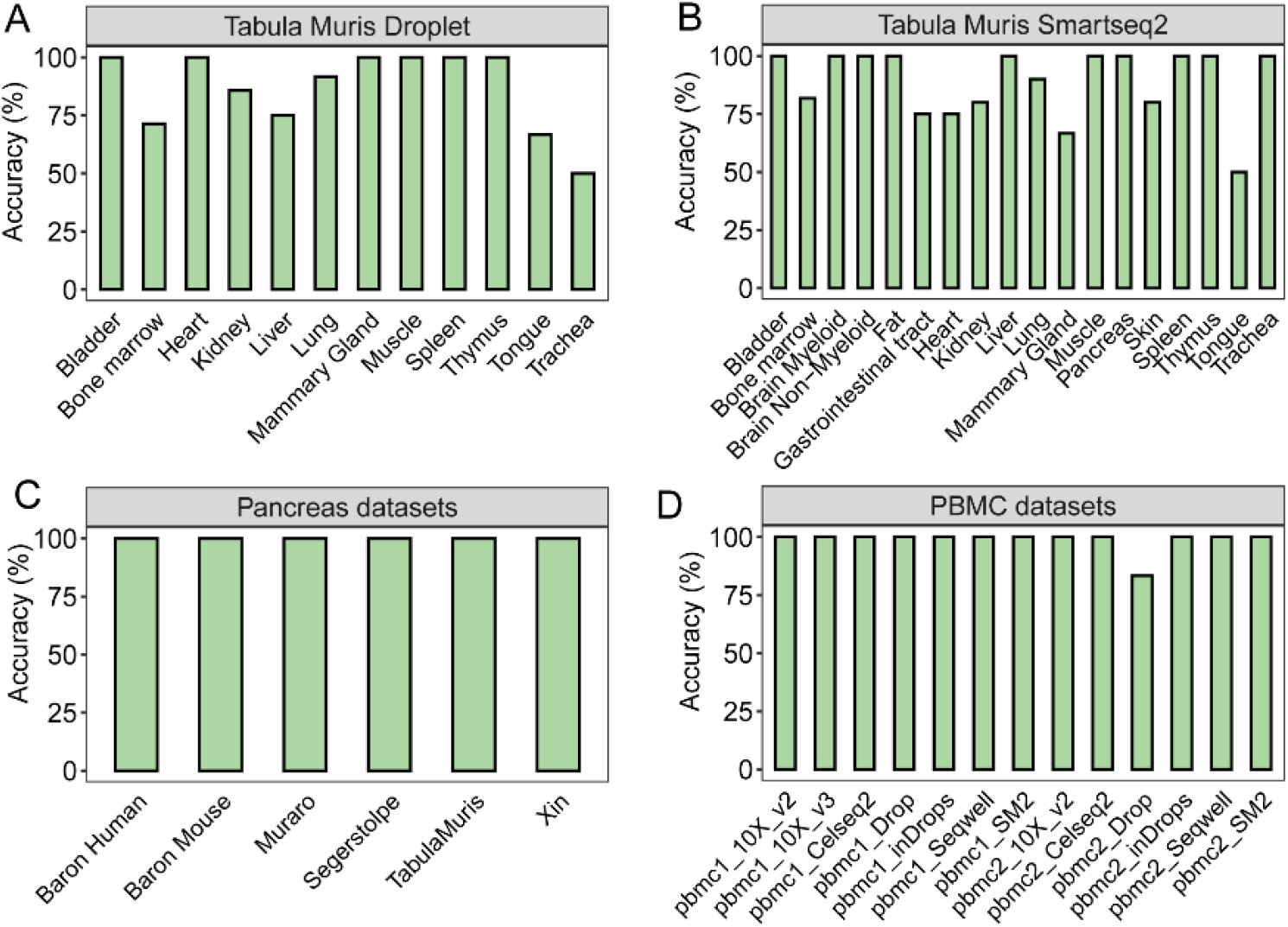
Performance of scMayoMap on experimental datasets cover different tissues. Barplot shows the annotation accuracy of scMayoMap across (A) 12 different tissues on the Tabula Muris Droplet scRNA-seq datasets, (B) 18 tissue types on the Tabula Muris Smartseq2 scRNA-seq datasets, (C) 6 different pancreas scRNA-seq datasets from literature, and (D) 13 different PBMC scRNA-seq datasets. Accuracy is calculated as percentage of clusters correctly annotated by each method.

A great amount of scRNA-seq data has been generated in peripheral blood mononuclear cells (PBMCs) because of the high interest and accessibility in both basic and clinical research. Next, we performed a second level of annotation to identify the subpopulations of immune cells in PBMCs, such as CD4 naïve T cells, CD4 memory T cells, CD8 naïve T cells, CD8 central memory T cells, CD8 effector memory T cells, etc. Within 12 of the 13 tested PBMC datasets, *scMayoMap* correctly identified all cell types with accurate annotation of subtypes in T cells and monocytes. Only one cluster was mis-annotated by *scMayoMap* in the pbmc2_Drop dataset (Fig. 3D and Table S4). A good annotation tool should be compatible with different scRNA-seq protocols. These 13 PBMC datasets were generated by 7 types of sequencing technologies, including two low-throughput plate-based methods (Smart-seq2 and CEL-Seq2) and five high-throughput methods (10x Chromium, Drop-seq, Seq-Well, and inDrops). *scMayoMap* performed well across all these different scRNA-seq platforms, indicating that it is highly accurate and applicable to a wide range of tissues using different scRNA-seq methods.

### 3.3 The performance of *scMayoMap* is superior to other annotation tools

In clinical studies, PBMCs are regularly obtained and furnish valuable information related to diseases and treatments. To compare the performance of *scMayoMap* with other available annotation tools, we firstly compared the performance of *scMayoMap* to *SingleR, SCSA, scCATCH*, and *scmap* on the 13 PBMC datasets. As a result, *scMayoMap* successfully identified all cell types in most of the datasets with a mean accuracy of 99% (range: 83%-100%, median: 100%), comparing to 93% for *SingleR*, 88% for *scmap*, 72% for *SCSA*, 51% for *scCATCH* (Fig. 4A, Table S4-S6).

**Figure 4.**
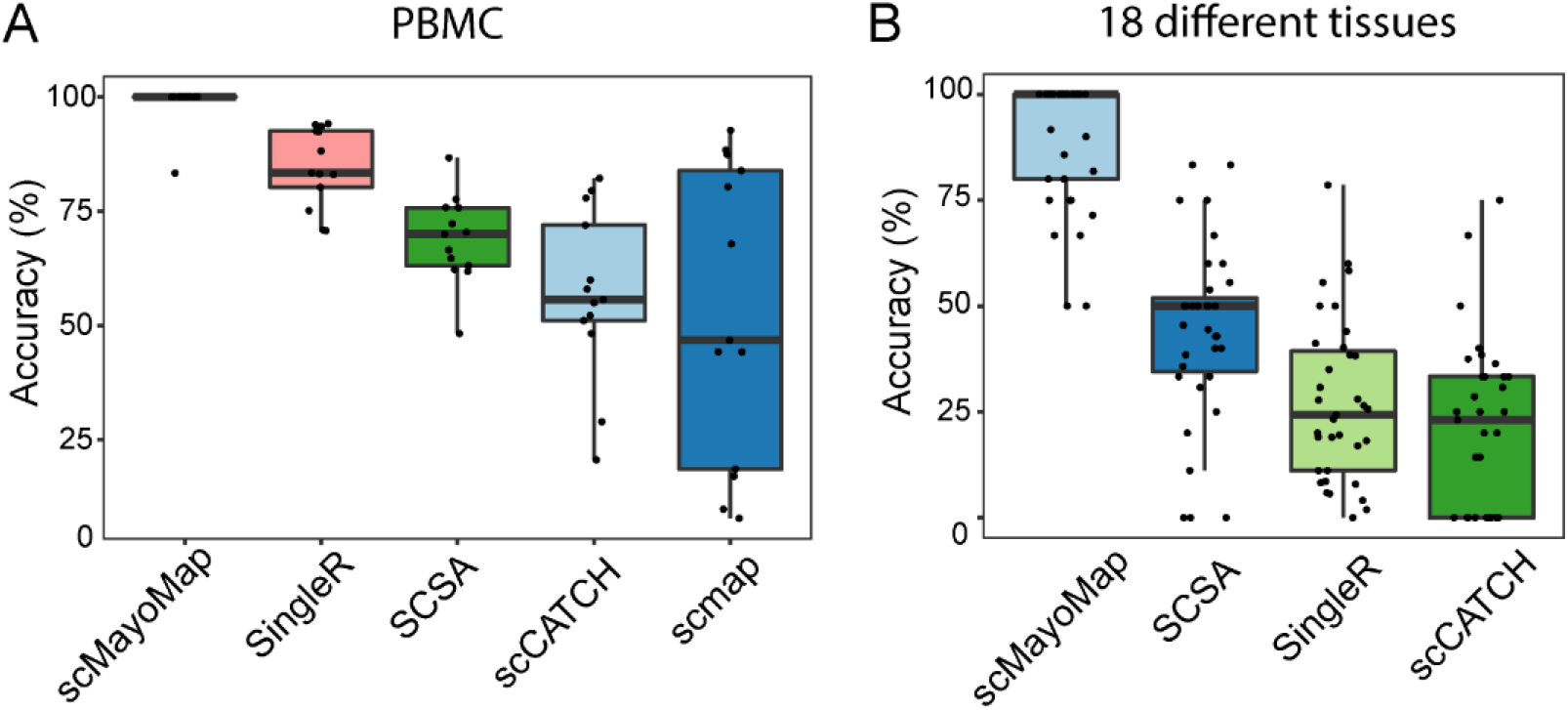
Comparison of scMayoMap and other cell annotation tools on 48 real datasets. (A) Comparison of celltype annotation accuracy in PBMC datasets. Thirteen PBMC scRNA-seq datasets generated by seven sequencing technologies were tested, including two low-throughput plate-based methods (Smart-seq2 and CEL-Seq2) and five high-throughput methods (10x Chromium, Drop-seq, Seq-Well, and inDrops). (B) Comparison of cell type annotation accuracy on 35 datasets with different tissues from Tabula Muris and literature. The accuracy of each method is presented as the percentage of cells that were correctly annotated by each method. Jitters on the plot represents datasets. The whisker extends from the hinge to the value that is within 1.5 * interquartile range (IQR) of the hinge, where IQR is the inter-quartile range, or distance between the first and third quartiles.

Further, we assessed the annotation accuracy of *scMayoMap* and *SingleR, SCSA, and scCATCH* on 35 datasets from 18 tissues of *Tabula Muris* datasets and five additional datasets from literature (Fig. 4B). The findings indicated that *scMayoMap* outperforms other methods, with a mean accuracy of 90% (50%-100%, median: 100%). In contrast, *SCSA* showed a mean accuracy of 44% (0%-83%, median: 50%), *SingleR* showed a mean accuracy of 27% (1%-79%, median: 24%), and *scCATCH* showed a mean accuracy of 22% (0%-75%, median: 23%).

Moreover, *scMayoMap* was used in multiple different tissues, including mouse skeletal muscle [27], brain [28], kidney [29], in datasets generated in our lab and provided perfect cell annotation accuracy. Overall, the accuracy of *scMayoMapDatabase* and *scMayoMap* is superior to the existing databases and tools.

### 3.4 *scMayoMapDatabase* can improve the annotation accuracy of other tools

We made *scMayoMapDatabase* easily accessible from the package and evaluated its performance as a replacement for the internal database of other annotation tools. To test this, we replaced the default database used in SCSA with scMayoMapDatabase due to its high flexibility. Interestingly, this led a substantial improvement in the performance of *SCSA*; the prediction accuracy of 9 out of 13 datasets increased (range: 3%-38%; median: 18%) (Fig. 5). These results suggest that *scMayoMapDatabase* contributes significantly to the high performance of scMayoMap, and it can be a valuable resource for cell type annotation in scRNA-seq analysis.

**Figure 5.**
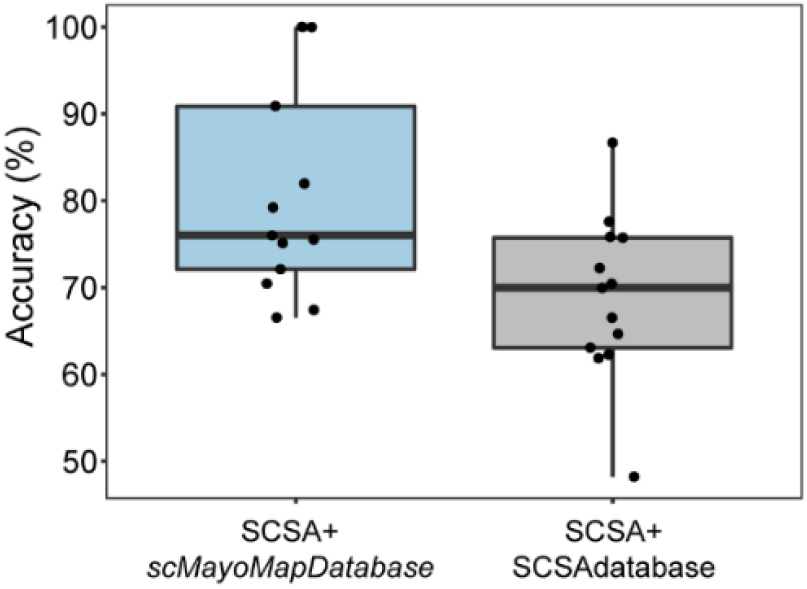
scMayoMapDatabase improves the annotation accuracy of SCSA. The cell types within the 13 PBMC datasets (same as in Figure 3D) were predicted by SCSA using its internal database or scMayoMapDatabase.

## 4. Discussion

We developed a new scRNA-seq data annotation tool, simplifying the cell type annotation process for scRNA-seq data. By using standard clustering results from upstream analysis as input, *scMayoMap* can swiftly annotate the corresponding cell types for each cluster, based on its internal cell marker database, *scMayoMapDatabase*. Through testing experimental datasets with varying cell numbers and types, we found that scMayoMap can complete annotation in under a minute by running a single function line included in the scMayoMap package. This enhanced usability of the tool is particularly beneficial for researchers, including those in clinical settings.

The reference database plays a critical role in cell type annotation tools. While several existing cell marker databases have helped advance research using scRNA-seq, they are not without limitations, such as inadequate or inappropriate cell markers and insufficient cell types or tissue coverage. Additionally, there are numerous marker genes in these databases that are identified only by their nicknames, which lack standardization. To overcome these shortfalls, we generated the *scMayoMapDatabase* by integrating marker genes from current popular databases and manually curating the tissues as well as the cell types.

This curated database serves as the foundation for accurate cell type prediction, as demonstrated in our results.

By extensive evaluation on experimental datasets using different scRNA-seq techniques, we demonstrated that the performance of *scMayoMap* surpassed other methods, including cluster annotation methods and the cell annotation methods. A previous study illustrated that current methods using prior knowledge of marker genes did not show promising performance on PBMC datasets [18]. In this study, we demonstrate that *scMayoMap* can predict PBMC cell types with small errors, suggesting that marker-based approach is still a promising approach if applied properly. Additionally, by extensive evaluation on experimental datasets of different tissues, we demonstrate that *scMayoMap* is a useful tool for cell type annotation with high accuracy.

We acknowledge that *scMayoMap* has certain limitations, such as its inability to identify new cell types since its predictions rely on an existing marker database. However, it can still provide insights for identifying novel cell types by returning the closest cell types and their corresponding marker genes. We also caution users to be careful when interpreting the prediction results of a cluster that has multiple cell types assigned to it. In order to assist this, we have incorporated an output of evaluation score for clusters with multiple cell types assigned. This score can be easily accessed using the “scMayoMap.obj$markers” command as explained in the package tutorial. Additionally, users can use their biological knowledge to analyze the expression of marker genes retrieved for each cluster to gain further insights.

Overall, *scMayoMap*, combined with the comprehensive scMayoMapDatabase, is a user-friendly and powerful tool for annotating cell types in scRNA-seq data analysis, with potential applications in scRNA-seq studies.

## Acknowledgments

We are grateful for the support of the Robert and Arlene Kogod Center on Aging, the Center for Individualized Medicine, and the Center for Biomedical Discovery at Mayo Clinic. We thank Drs. Marissa Schafer, Yi Zhu, Aleksey Matveyenko, Jielu Hao, Madison Doolittle, Chunhua Chen, Bin Zhang, and Hong Cao for reviewing the *scMayoMapDatabase*. We thank Dr. Ming Xu and Rachel Cohn for testing the package. Figure 1A was generated in Biorender.

## Availability of Data and Materials

*scMayoMap* is an open-source software, available at GitHub (https://github.com/chloelulu/scMayoMap). The *scMayoMapDatabase* and raw data for each figure can be found in the supplementary table. Experimental datasets analyzed and code used in this study can be found at: https://github.com/chloelulu/scMayoMap.

